# The impact of tropodithietic acid on microbial physiology under varying culture complexities

**DOI:** 10.1101/2024.10.23.619853

**Authors:** Olesia Shlakhter, Einat Segev

## Abstract

Research has increasingly focused on understanding marine bacterial physiology under environmentally relevant conditions. Biotic interactions have a key influence on microbial physiology and are often modeled in the lab by manipulating the complexity of microbial cultures. Notably, findings from low-complexity cultures that have been re-evaluated in more complex systems, occasionally lead to different outcomes. Here, we assess how the genomic capability of bacteria to produce secondary metabolites, specifically the antibiotic tropodithietic acid (TDA), influences microbial physiology and interactions, under varying microbial complexity.

We investigate the impact of TDA production on microbial physiology across systems with increasing complexity: from bacterial mono-cultures, to bacterial co-cultures of *Phaeobacter inhibens* with *Dinoroseobacter shibae*, and a more complex tri-culture that includes both bacteria and their algal host *Emiliania huxleyi*. In these systems, we examine both wild-type (WT) *P. inhibens* bacteria and mutant bacteria with a *tdaB* deletion (Δ*tdaB*) that are not capable of producing TDA. This systematic approach allowed us to explore the relationship between the *tdaB* gene, microbial physiology, and system complexity.

Our findings show that deleting the *tdaB* gene affected bacteria-bacteria interactions in co-cultures but not in tri-cultures with the algal host. Additionally, our data revealed that algal death was delayed in cultures containing *P. inhibens* Δ*tdaB* mutants compared to those with WT bacteria.

Results of our study highlight the importance of microbial complexity in the study of bacterial physiology and point to the understudied role of TDA in algal-bacterial interactions.

**Importance:** This study advances our understanding of marine bacterial physiology under varying levels of microbial complexity. We uncover how the production of secondary metabolites, specifically the antibiotic tropodithietic acid (TDA), influences microbial interactions. Our systematic approach, which includes bacterial mono-cultures, co-cultures, and tri-cultures involving algal hosts, allows us to evaluate the impact of the *tdaB* gene on microbial interactions. Notably, our findings reveal that the deletion of this gene affects bacteria-bacteria interactions in co-culture but does not have the same effect in more complex systems that include algae. Additionally, the observation that algal death is delayed in cultures with TDA-deficient bacteria underscores the significance of TDA in these interactions. Overall, our research highlights the influence of the microbial culture complexity on bacterial physiology and emphasizes the overlooked role of TDA in algal-bacterial dynamics.

## Introduction

In recent years, research has increasingly concentrated on understanding marine bacterial physiology under environmentally relevant conditions. One key factor influencing microbial physiology is biotic interactions, which are often simulated in the lab through the microbial complexity of the cultures. For instance, bacterial physiology has been shown to be affected by various algal metabolites, as evidenced by studies on algal-bacterial cultures. Algal-driven bacterial responses include accelerated bacterial growth (1), formation of a bacterial extracellular matrix (2), enhanced bacterial conjugation (3), and increased synthesis of secondary metabolites (4). Additionally, bacteria grown in the presence of other bacterial species have demonstrated altered physiology, such as the production of novel secondary metabolites (5) or increased antibiotic production (6).

Notably, findings from cultures with low complexity that have been reassessed in more complex systems, occasionally revealed different outcomes. For example, the pathogenicity of *Phaeobacter inhibens* bacteria towards *Emiliania huxleyi* algae was initially observed in algal-bacterial co-cultures (7–10). However, when this pathogenic behavior was examined in a community with additional bacteria, it was found that the pathogenicity did not manifest (11). It was also demonstrated that the effect of *P. inhibens* on host-associated microbiomes varies depending on the complexity and composition of the existing microbiome (12). Therefore, understanding microbial physiology in laboratory models should account for the impact of microbial complexity within the system.

Here, we aimed to evaluate how the bacterial capability to encode the synthesis of a secondary metabolite influences microbial physiology, under varying microbial complexities.

*P. inhibens* bacteria produce the secondary metabolite tropodithietic acid (TDA) which acts as an antibiotic against various organisms such as the fish pathogenic bacteria of the species *Vibrio anguillarum* (13, 14). The synthesis of TDA can be completely abolished by deletion of the *tdaB* gene (15). *P. inhibens* bacteria are commonly found in algal blooms, where they interact with various co-occurring bacteria as well as with their algal hosts (8, 10). To study how the ability of *P. inhibens* to produce TDA affects microbial physiology within different contexts of microbial complexity, we established both a co-cultivation system of *P. inhibens* with *Dinoroseobacter shibae* bacteria and a tri-cultivation system with the addition of the *Emiliania huxleyi* algal host. We incorporated either wild-type (WT) *P. inhibens* or *tdaB-*deleted mutants (*ΔtdaB*) into these co- and tri-cultures to assess the relationship between the *tdaB* gene, microbial physiology, and microbial complexity. Our findings reveal the importance of microbial complexity in the study of bacterial physiology and highlight the role of TDA in both bacterial-bacterial and algal-bacterial interactions.

## Results

### Establishing experimental systems to study the impact of TDA on microbial interactions

To investigate the impact of TDA on the interactions between *P. inhibens* and other microorganisms, we established three experimental systems with varying levels of microbial complexity (see Fig. 1). Each system utilized either wild-type (WT) *P. inhibens* or mutant *P. inhibens* lacking the *tdaB* gene (*ΔtdaB*), which is essential for TDA production.

**Figure 1.**
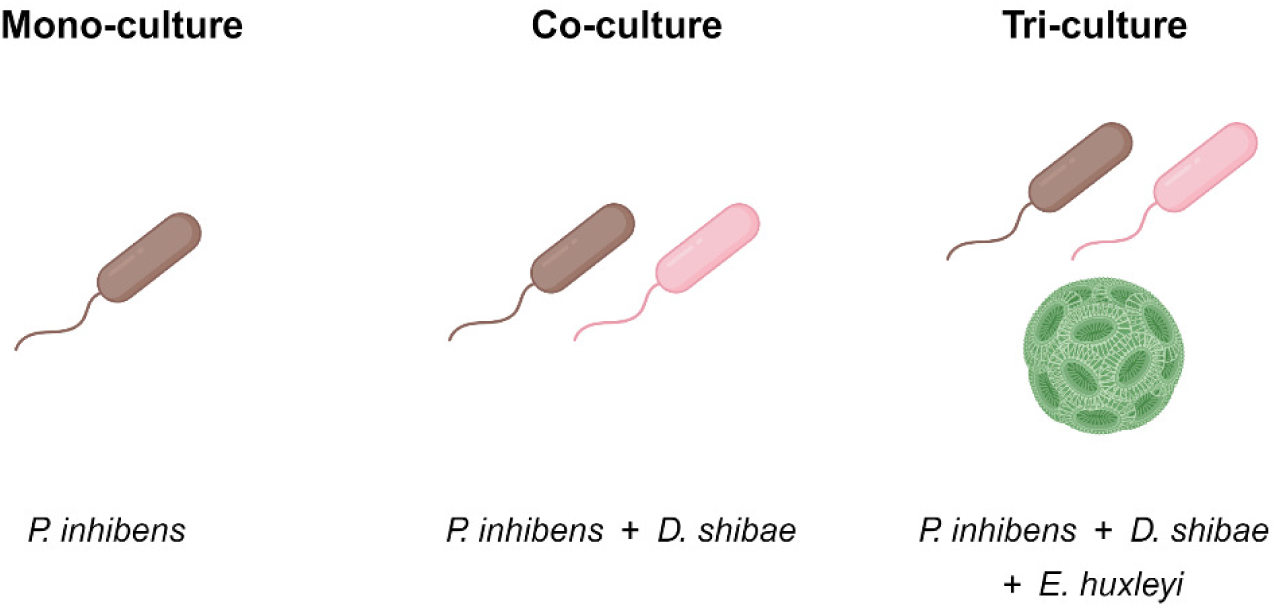
Schematic representation of the microbial cultures used in this study. Mono-cultures contain pure cultures of *P. inhibens* DSM 17395 wild-type (WT) or Δ*tdaB* mutant bacteria (as indicated), in CNS medium (see Materials and Methods) with 5.5 mM succinate as a sole carbon source. Co-cultures contain *P. inhibens* bacteria (WT or Δ*tdaB,* as indicated) with *D. shibae* DFL-12 bacteria in CNS medium containing 5.5 mM succinate as a sole carbon source. Tri-cultures contain *P. inhibens* bacteria (WT or Δ*tdaB,* as indicated), *D. shibae* bacteria, and *E. huxleyi* algae in synthetic seawater (see materials and methods) with no added carbon source.

The simplest system was a mono-culture of either *P. inhibens* WT or *ΔtdaB*, grown in a defined medium with succinate as the sole carbon source (Fig. 1A). Mono-culture cell counts over time were achieved by monitoring OD_600_ (see Materials and Methods).

To increase microbial complexity, we created bacterial co-cultures of *P. inhibens* (WT or *ΔtdaB*) with *Dinoroseobacter shibae* (Fig. 1B). *D. shibae* bacteria naturally coexists with *P. inhibens* bacteria during algal blooms of the microalga *Emiliania huxleyi* (10). In this co-culture system, bacteria were cultivated in a defined medium with succinate as the sole carbon source. To accurately enumerate bacterial cells of the two species, we used selective plates to distinguish between them (see Materials and Methods). The selective plates allowed precise colony-forming unit (CFU) counts for each species, facilitating clear differentiation and quantification (see Materials and Methods).

The most complex system in our study was a tri-culture, which combined both bacterial species (*P. inhibens* WT or Δ*tdaB* and *D. shibae*) with their algal host, *E. huxleyi* (Fig. 1C). These tri-cultures were grown in artificial seawater without any additional external carbon sources, compelling bacteria to depend on organic carbon from algal exudates.

In tri-cultures, bacterial cell enumeration was achieved using CFU counts on selective plates, complemented with a second independent method of quantitative PCR (qPCR) to validate the results (see Materials and Methods).

The growth curves of mono-cultures were comparable between the WT and Δ*tdaB* strains of *P. inhibens*. Previous studies demonstrated that mutant strains with deletions in genes encoding secondary metabolites can exhibit increased bacterial growth, highlighting the potential impact of such genetic modifications (16). However, we observed only minor differences in bacterial growth between WT and Δ*tdaB P. inhibens* bacteria in mono-cultures (Fig. 2). The comparable growth of WT and mutant *P. inhibens* strains in mono-culture provides a baseline for assessing the impact of TDA on microbial interactions; Any differences in growth observed in the presence of other bacteria or algae can highlight the possible influence of TDA on these interactions.

**Figure 2.**
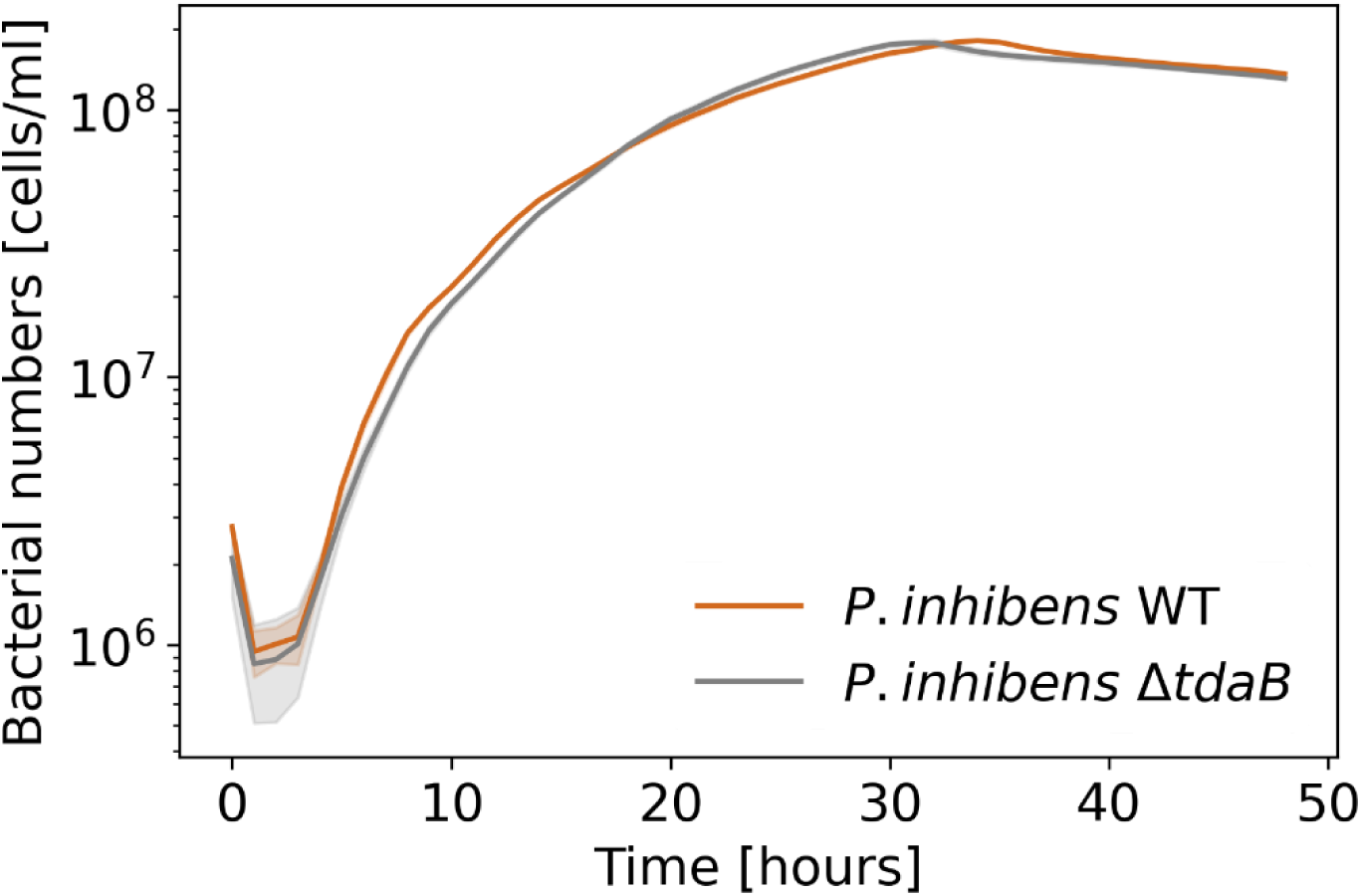
Deletion of the *tdaB* gene does not affect the growth of *P. inhibens* bacteria in mono-culture. Growth of *P. inhibens* WT (brown) and Δ*tdaB* mutant (grey) bacteria in CNS medium containing succinate as a sole carbon source. Bacterial growth was monitored using OD_600_ and values were converted to cells/ml (see Materials and Methods). Each line represents the mean of four biological replicates, and shaded areas indicate the standard deviation.

### Deletion of the *tdaB* gene impacts the fitness of *P. inhibens* in co-cultures with *D. shibae*

To evaluate the impact of TDA on the interaction of *P. inhibens* and *D. shibae* bacteria, we first tested the susceptibility of *D. shibae* to TDA. We therefore plated a *D. shibae* lawn on Marine Broth (MB) agar plates and spotted *P. inhibens* WT and Δ*tdaB* cultures on top of the lawn. If *D. shibae* bacteria are killed by TDA, an inhibition zone should be visible around colonies of WT *P. inhibens* bacteria but not around colonies of the Δ*tdaB* strain. Our results show that WT *P. inhibens* colonies exhibit the typical brown pigmentation associated with TDA production (15, 17) and an inhibition zone around the colonies was evident (Fig. 3A, top). In contrast, colonies of Δ*tdaB P. inhibens* lost their pigmentation as previously described (15) and had no visible inhibition zone (Fig. 3A, bottom). These results confirmed that *D. shibae* is indeed susceptible to TDA produced by *P. inhibens*, and that the Δ*tdaB P. inhibens* strain is perturbed in TDA production.

**Figure 3.**
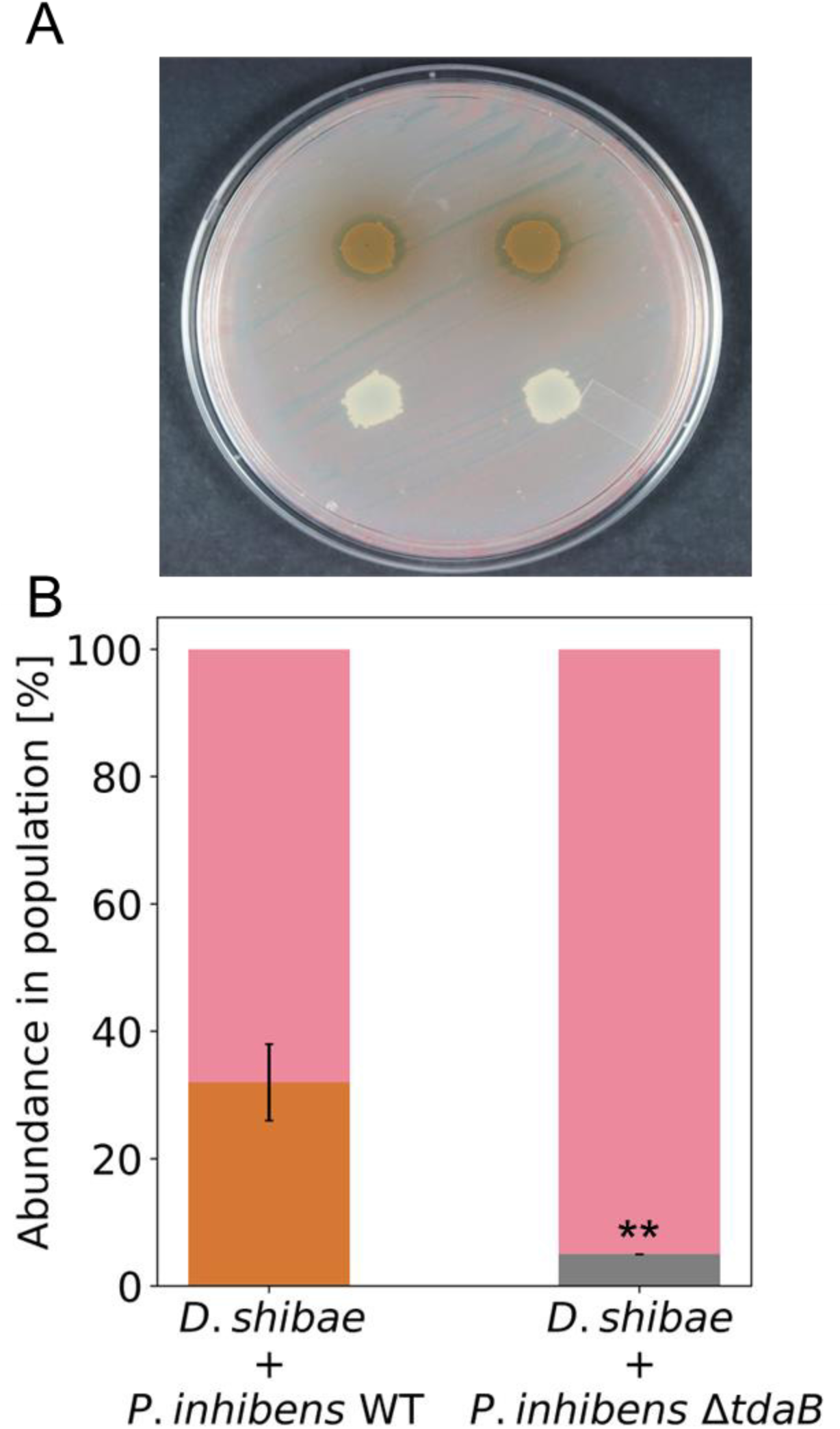
Deletion of the *tdaB* gene decreases the fitness of *P. inhibens bacteria* in co-cultures with *D. shibae*. **A** A lawn of *D. shibae* was plated on a MB plate and spotted with 5 ml of *P. inhibens* WT (upper, brown colonies) and Δ*tdaB* mutant (lower, white colonies) cultures. Plates were incubated for 72 hours at 30°C. Deletion of the *tdaB* gene resulted in loss of pigmentation as previously described (15) and no visible inhibition zone, characteristic of tropodithietic acid (TDA) secretion (17). **B** Relative abundance of *P. inhibens* WT (brown) and Δ*tdaB* (grey) bacteria, and *D. shibae* bacteria (pink) after 24 hours in co-culture with CNS medium containing 5.5 mM succinate as a sole carbon source. The initial inoculum was 1.5 x 10^6^ cells/ml for each bacterial strain (see Materials and Methods). Bars represent the mean value of three biological replicates, and error bars correspond to the standard deviation. \*\**p*<0.005 using a two-tailed t-test.

Next, we examined whether TDA production affects bacterial fitness in bacterial co-cultures. In liquid cultures, secondary metabolites (including TDA) are often produced by bacteria during later stages of growth as a result of nutrient limitations (18). Therefore, bacterial abundance in co-cultures was examined after 24 hours of growth, which corresponds to the transition between the late exponential and early stationary growth phases for both bacterial species (Fig. 2, Supplementary Fig. 1). Our results indicate that *P. inhibens* Δ*tdaB* bacteria exhibit decreased fitness compared to WT bacteria in co-cultures with *D. shibae* (Fig. 3B). Although deletion of the *tdaB* gene had no measurable effect on bacterial growth in mono-cultures, this mutation reduced the growth of *P. inhibens* bacteria in a co-culture, without affecting the growth of *D. shibae* bacteria (Table 1).

**Table 1:** The decreased fitness of *P. inhibens* Δ*tdaB* in co-cultures is the result of reduced growth of the mutants and not increased growth of *D. shibae*. Bacterial numbers in co-cultures were monitored using selective plates following 24 hours of growth in CNS medium containing 5.5 mM succinate as a sole carbon source. The initial inoculum of each bacterial strain was 1.5 x 10^6^ cells/ml (data corresponds to Fig. 3B). A, B and C designate biological replicates. \*\**p*<0.005 using a two-tailed t-test.

| | Bacterial numbers [CFU/ml] $\times 10^6$ | | | |
| --- | --- | --- | --- | --- |
| | A | B | C | Mean ± SD |
| *P. inhibens* WT | 19 | 27 | 21 | 22±4 |
| *P. inhibens* $\Delta tdaB^{**}$ | 3 | 3 | 2 | 2±0.3 |
| *D. shibae* (+ WT *P. inhibens*) | 50 | 70 | 30 | 50±20 |
| *D. shibae* (+ $\Delta tdaB$ *P. inhibens*) | 48 | 51 | 45 | 48±3 |

### Deletion of the *tdaB* gene does not affect the fitness of *P. inhibens* and *D. shibae* bacteria in tri-cultures

In the marine environment, various heterotrophic bacteria rely on their algal hosts as a source of organic carbon (19, 20). Given the impact of *tdaB* deletion on the fitness of *P. inhibens* in bacterial co-cultures with *D. shibae* when succinate is provided as the carbon source, we aimed to determine whether this decreased fitness is also evident when *P. inhibens* and *D. shibae* rely on their algal host.

Our data show that in tri-culture, bacterial growth of both *P. inhibens* and *D. shibae* was not affected by *tdaB* gene deletion (Fig. 4A and B). We observed no significant differences in the growth of WT and Δ*tdaB P. inhibens* bacteria in tri-cultures, nor in the growth of *D. shibae* bacteria. To validate these results, we used another independent method to assess bacterial abundance, using qPCR to quantify DNA copy numbers on day 8 of cultivation. Results from both the selective plates and the qPCR were consistent, showing no significant differences in the growth of WT and *ΔtdaB P. inhibens* or *D. shibae* bacteria in tri-cultures (Supplementary Fig. 3). These results align with previous studies that demonstrated only a minor impact of TDA on the bacterial partners of diatom microalgae (21).

**Figure 4.**
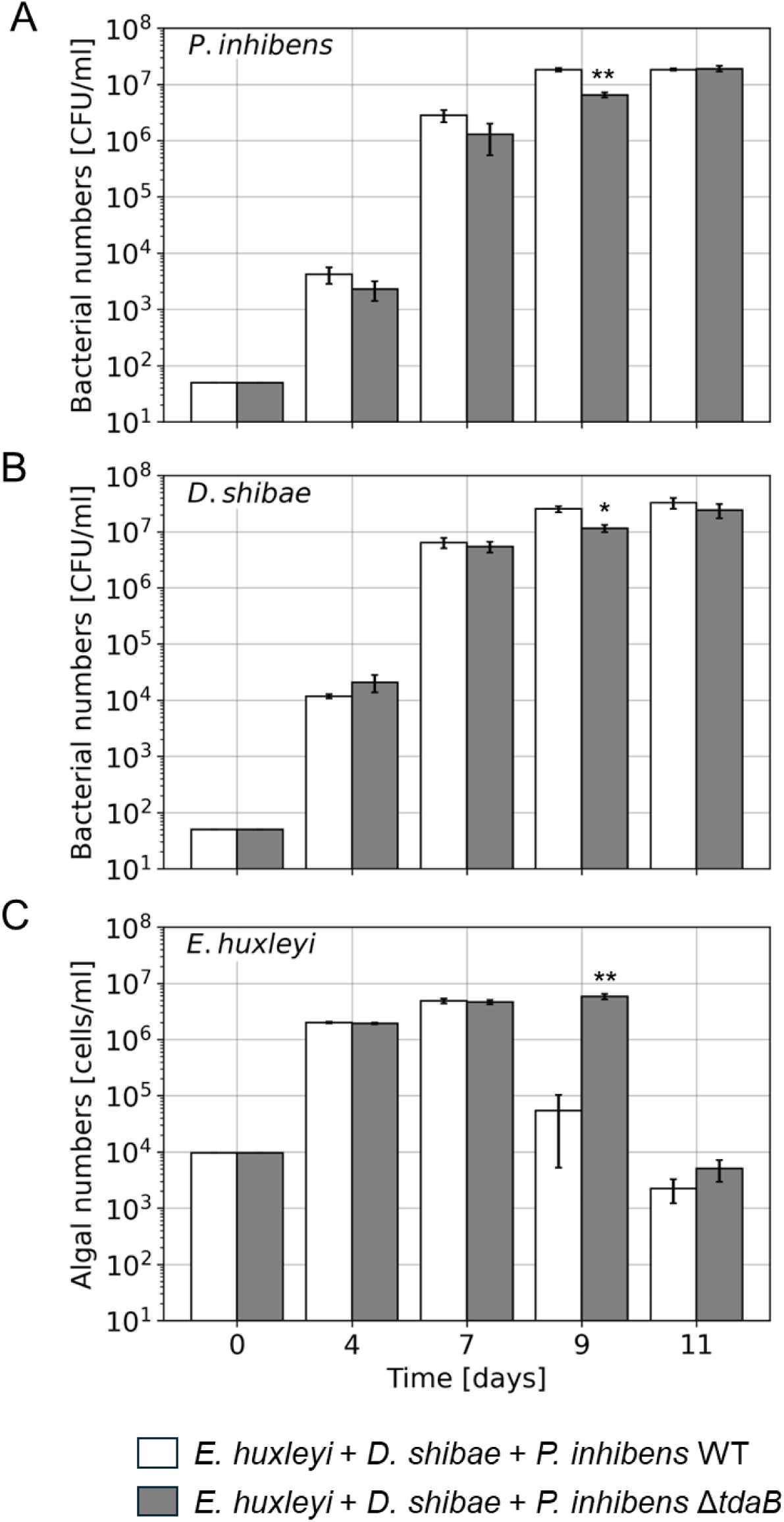
The deletion of the *tdaB* gene delays the pathogenicity of *P. inhibens* towards *E. huxleyi* algae in the context of tri-cultures. **A** Growth of *P. inhibens* WT bacteria (white bars) and Δ*tdaB* mutant bacteria (grey bars) in tri-cultures. **B** Growth of *D. shibae* bacteria in tri-cultures with *P. inhibens* WT strain (white bars) or the Δ*tdaB* mutant strain (grey bars). **C** Growth of *E. huxleyi* algae in tri-cultures with *D. shibae* bacteria and *P. inhibens* WT strain (white bars) or the Δ*tdaB* mutant strain (grey bars). Bacterial growth was monitored using selective plates, and algal growth was tracked using flow cytometry (see Materials and Methods). The differences in growth between cultures were evident at day 9 for all species. Bars represent mean values of three biological replicates, error bars indicate standard errors. * indicates *p*<0.05, ** indicates *p*<0.005 using a two-tailed t-test.

However, an interesting exception was noted on day 9; both *P. inhibens* and *D. shibae* showed decreased growth in tri-cultures containing the *ΔtdaB* strain. Since both bacterial species were similarly affected, it is difficult to attribute this decrease to a growth disadvantage specific to the absence of TDA. Instead, as discussed below, the reduced growth of both *P. inhibens* and *D. shibae* appears to reflect changes in the algal host population dynamics.

### Deletion of the *tdaB* gene affects the pathogenicity of *P. inhibens* towards the algal host

*P. inhibens* bacteria display pathogenic behavior towards *E. huxleyi* algae, leading to algal cell death (9, 10). Our data show that deletion of the *tdaB* gene resulted in delayed pathogenicity of *P. inhibens* towards *E. huxleyi* both in tri-cultures (Fig. 4C) and in cultures of only *P. inhibens* with *E. huxleyi* (Supplementary Fig. 2). This delay suggests that TDA production may be linked to the pathogenicity of *P. inhibens* and could influence algal-bacterial interactions. Although the exact relationship between bacterial TDA production and algal death is not yet fully understood, the lack of TDA production seems to delay algal death thereby affecting the availability of algal-derived organic carbon for the associated bacteria. In tri-cultures with WT *P. inhibens*, bacterial growth of both *P. inhibens* and *D. shibae* exceeds densities of 10^7^ cells/ml upon algal death. In contrast, when algal death is delayed in tri-cultures with the *ΔtdaB* strain, bacterial growth of both *P. inhibens* and *D. shibae* remains below 10^7^ cells/ml, and only following the delayed algal death, the density of the cultures exceeds 10^7^ cells/ml. In conclusion, TDA appears to play an overlooked role in both bacterial-bacterial and algal-bacterial interactions.

## Discussion

### The importance of culture complexity in laboratory studies of microbial physiology

Secondary metabolites, especially antibiotics, shape microbial interactions (22). In the marine environment, host-associated bacteria produce a wide range of antibiotics, as demonstrated by bacteria from the algal microbiome (23) and sponge microbiome (24, 25). In turn, antibiotics production is significantly impacted by interactions among bacteria. For example, marine bacteria from the *Streptomyces tenjimariensis* species produce the antibiotic istamycin to outcompete susceptible bacterial species (26). Bacterial interactions can also inhibit antibiotics production, as reported for the coral-associated actinobacterium *Agroccocus* that impaired the ability of another *Streptomyces* actinobacterium to produce antibiotics (27).

Therefore, when studying the impact of secondary metabolites on microbial physiology in laboratory model systems, it is important to consider the potential influence of the microbial complexity within the system. In this study, we examined the impact of deleting the *tdaB* gene, which is essential for the production of the TDA antibiotic in *P. inhibens*, on interactions with ecologically relevant partners. Cultures with varying levels of microbial complexity - from mono-cultures of *P. inhibens*, to co-cultures with *D. shibae* bacteria and tri-cultures with both bacteria and *E. huxleyi* algae - allowed us to evaluate the influence of the *tdaB* deletion on these interactions.

### The ability to produce a metabolite versus actual metabolite production

Deletion of the *tdaB* gene eliminates the bacterial ability to produce TDA (15), since the deletion of a key gene from a biosynthetic gene cluster typically leads to the loss of the ability to produce that specific metabolite. However, the presence of the *tdaB* gene does not guarantee the production of this metabolite, as many bacterial secondary metabolites genes were shown to be silent or cryptic in laboratory culture conditions (28). Furthermore, although all the necessary genes are present, antibiotic synthesis may sometimes require induction by other species (29–31).

Here, the deletion of the *tdaB* gene in *P. inhibens* resulted in loss of pigmentation and ability to create an inhibition zone (Fig. 3A), two phenotypes that are largely attributed to TDA production. Therefore, it is likely that differences in the growth dynamics observed between the WT and Δ*tdaB* cultures are the result of difference in TDA production. This interpretation is further supported by the observation that the mutation itself did not affect bacterial growth dynamics, as evident by bacterial growth in mono-cultures (Fig. 2).

### The impact of TDA on bacterial interactions depends on the culture complexity

The influence of the *tdaB* gene deletion on bacterial interactions appears to depend on the culture complexity. In our study, deleting the *tdaB* gene in *P. inhibens* affected its interaction with *D. shibae* in co-culture (Fig. 3), but not in tri-culture when the algal host *E. huxleyi* was present (Fig. 4). A key difference between co-cultures and tri-cultures that should be noted, is the composition and flux of the bacterial nutrients supply. In co-cultures, bacteria receive a defined medium with succinate as a sole carbon source. Essential nutrients are supplied as fresh medium at the onset of the experiment and are gradually depleted as bacteria consume them during growth. Under these conditions, competition between *P. inhibens* and *D. shibae* is likely to occur, consequently driving the expression of the *P. inhibens tdaB* to inhibit *D. shibae*. However, in tri-cultures with the algal host, bacteria experience a continuous flux of algal secreted compounds-termed algal exudates-consisting of a complex mixture of nutrients and biomolecules (32, 33). The variety of compounds in algal exudates can allow different bacteria to consume different resources, thereby driving niche partitioning and preventing competition (34–36). Moreover, algal exudates can impact the expression of various biosynthetic pathways in *P. inhibens* (4, 10, 37). Therefore, it is possible that *tdaB* expression is regulated differently in co-cultures versus tri-cultures, resulting in different levels of TDA. Interestingly, a previous study demonstrated that *P. inhibens* bacteria actively expresses *tdaB* and *tdaC* genes when cultivated with algal hosts (38). Moreover, *P. inhibens* WT but not TDA-negative mutants were shown to decrease the concentration of the pathogen *Vibrio anguillarum* when cultivated along with an algal host (39). TDA was also suggested to contribute to the ability of *P. inhibens* bacteria to decrease the relative abundance of pathogenic *Vibrionales* in the oyster microbiome (12).

### A potential link between TDA and bacterial pathogenicity

Bacterial antibiotics are widely studied in the context of bacterial interactions (40, 41) but their role in algal-bacterial associations is less understood. In the current study, the ability of *P. inhibens* to produce TDA had an impact on the algal host. *P. inhibens* WT bacteria induce algal death, a phenomenon that was previously studied by us (7, 10, 11) and by others (8, 9, 42). Our data now show that algal death was delayed in cultures with *P. inhibens* Δ*tdaB* mutants compared to WT bacteria. The delay in algal death occurred in both tri-cultures (Fig. 4) and in cultures with *E. huxleyi* and *P. inhibens* Δ*tdaB* (Supplementary Fig. 2). Whether TDA is directly involved in bacterial pathogenicity, or whether the TDA impact on bacterial physiology has an indirect influence on pathogenicity, is yet to be understood.

Interestingly, various iron-binding molecules such as siderophores (4, 43) and roseochelins (44) are known bacterial virulence factors that are related to bacterial pathogenicity towards the host. Previous studies demonstrated that iron is also required for the production of active TDA (17). Therefore, iron emerges as a micronutrient that merits further study in the context of antibiotics-mediated algal-bacterial interactions.

To conclude, our study demonstrates the importance of microbial complexity in the study of bacterial physiology and highlights gaps in our understanding of the various roles that secondary metabolites play in microbial interactions.

## Materials and Methods

### Strains and growth conditions

The algal axenic strain *Emiliania huxleyi* CCMP3266 was purchased from the National Center for Marine Algae and Microbiota (Bigelow Laboratory for Ocean Sciences, Maine, USA). Algae were cultivated in Artificial Sea Water (ASW) containing L1-Si supplements (Na_2_SiO_3_-omitted L1 medium according to Guillard and Hargraves (45)). Cultures were kept in 18°C under a light/dark cycle of 16/8 hours and illumination intensity of 130 μmol photons m^−2^ s^−1^. The axenic state of algal cultures was monitored weekly by plating and under the microscope.

The bacterial strain *Phaeobacter inhibens* DSM 17395 was purchased from the German Collection of Microorganism and Cell Cultures (DSMZ, Braunschweig, Germany). Bacteria were plated from glycerol stocks (stored at -80°C) on ½ YTSS agar plates (2 g yeast extract, Sigma-Aldrich, USA; 1.25 g tryptone, Sigma-Aldrich, USA; 20 g sea salts, Sigma-Aldrich, USA; and 16 g agar per liter, BD Difco, USA), and after 48 hours of incubation at 30°C, single colonies were inoculated into liquid CNS medium for a pre-culture (5.5 mM glucose, Sigma-Aldrich, USA; 5 mM NH_4_Cl, Sigma-Aldrich, USA; 33 mM NaSO_4_, Fisher Scientific, UK; and diluted in ASW). Pre-cultures were cultivated in 30°C in the dark with 130 rpm shaking for at least 48 hours to reach the stationary phase.

The bacterial strain *Dinoroseobater shibae* DFL-12 (DSM 16493) was received from the lab of Prof. Irene Wagner Dobler (DSMZ, Germany). Bacteria were plated from glycerol stocks on MB agar plates (37.4 g MB powder, BD Difco, USA; and 16 g agar per liter, BD Difco, USA). Single colonies were inoculated into liquid CNS medium for a pre-culture (5.5 mM sodium succinate, Sigma-Aldrich, USA; 5 mM NH_4_Cl, 33 mM NaSO_4_, and diluted in ASW) and cultivated in 30°C in the dark with 130 rpm shaking for at least 48 hours.

The mutant strain *Phaeobacter inhibens* DSM17395 *ΔtdaB* was generated in the current study (see below) and cultivated under the same conditions used for the wild-type (WT) *P. inhibens* DSM17395 strain, with the addition of 30 μg/ml gentamicin.

### Deletion of the *tdaB* gene in *P. inhibens*

To replace the *tdaB* gene of *P. inhibens* DSM 17395 with an antibiotic resistance gene, the mutant strain ES150 was generated by inserting a gentamicin-resistance gene into the gene PGA1_262p00970. Regions of approximately 1000 bp upstream and downstream of the gene were PCR-amplified using primers 1021, 1022, 1023 and 1024, respectively (Table 2). The amplified fragments were assembled and cloned into the pDN18 vector (Table 3) using restriction-free cloning (46, 47). The resulting plasmid pES1 (Table 3) was introduced into competent *P. inhibens* bacteria by electroporation.

**Table 2:**
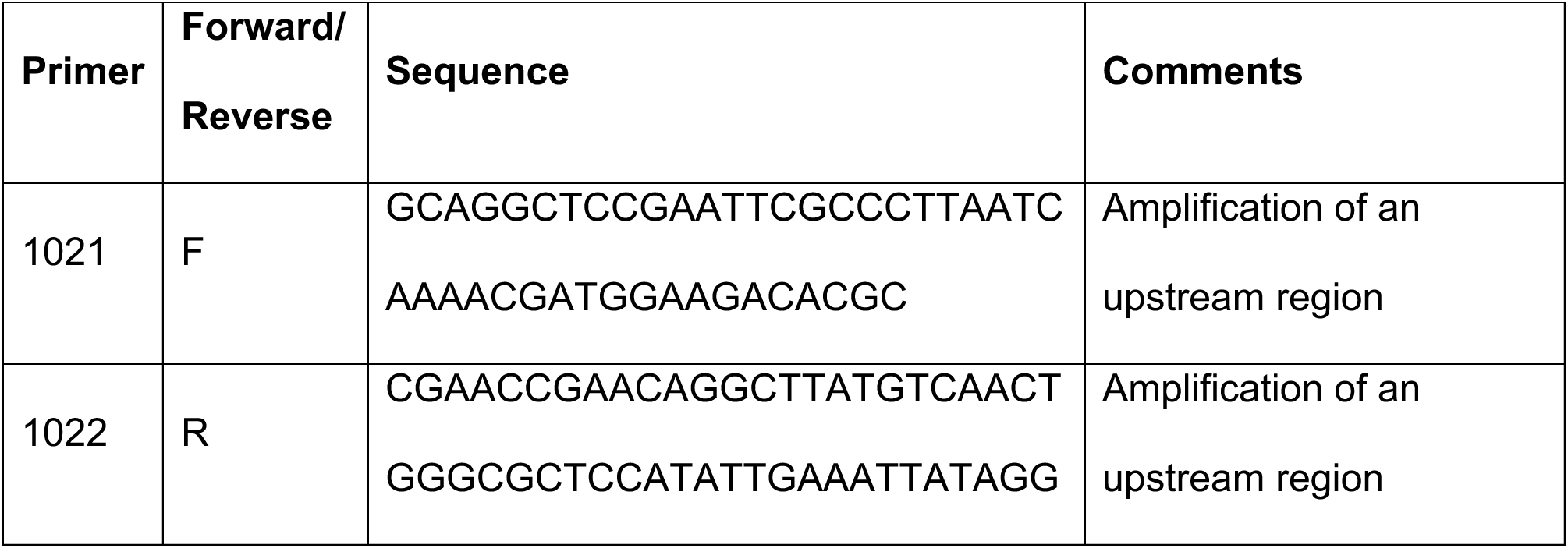
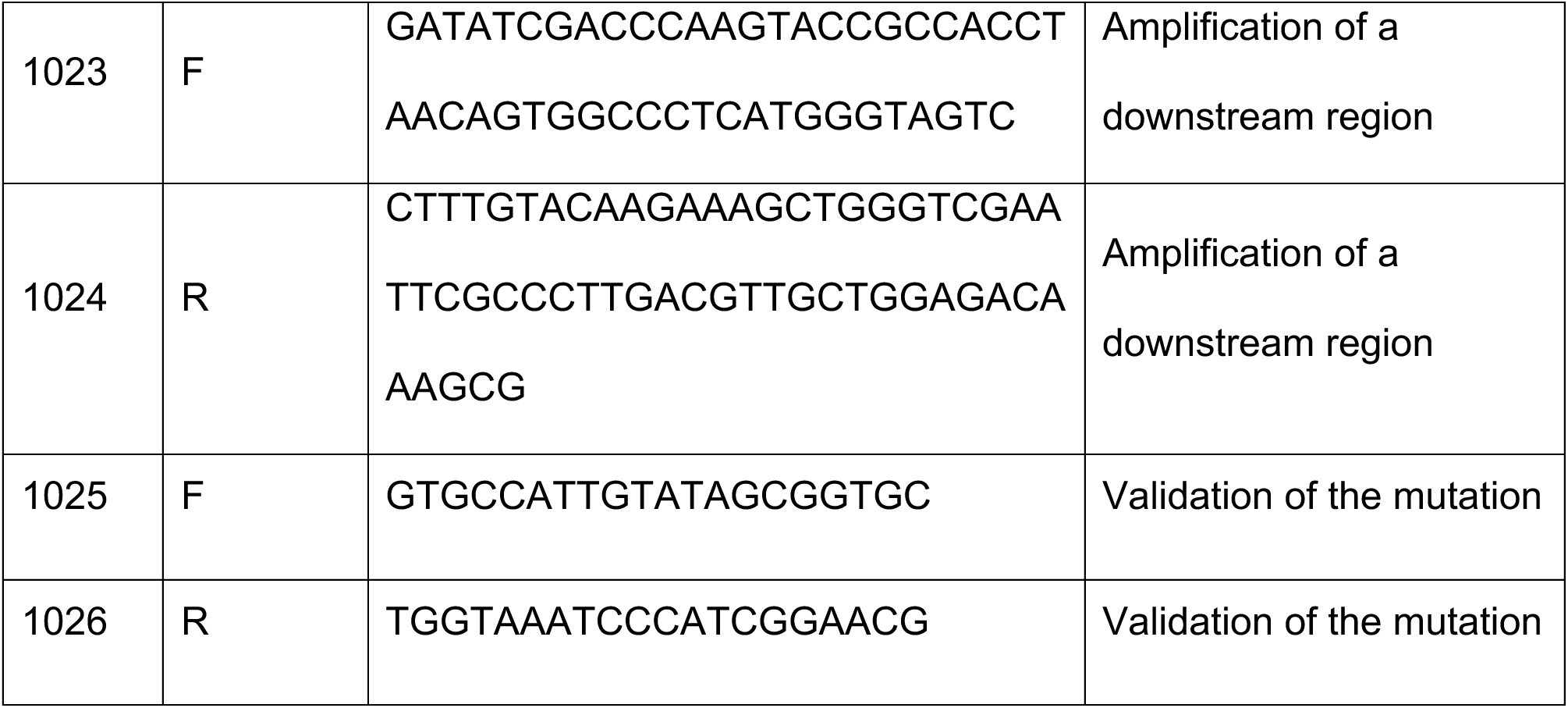
Primers used in the study for mutagenesis.

**Table 3:**
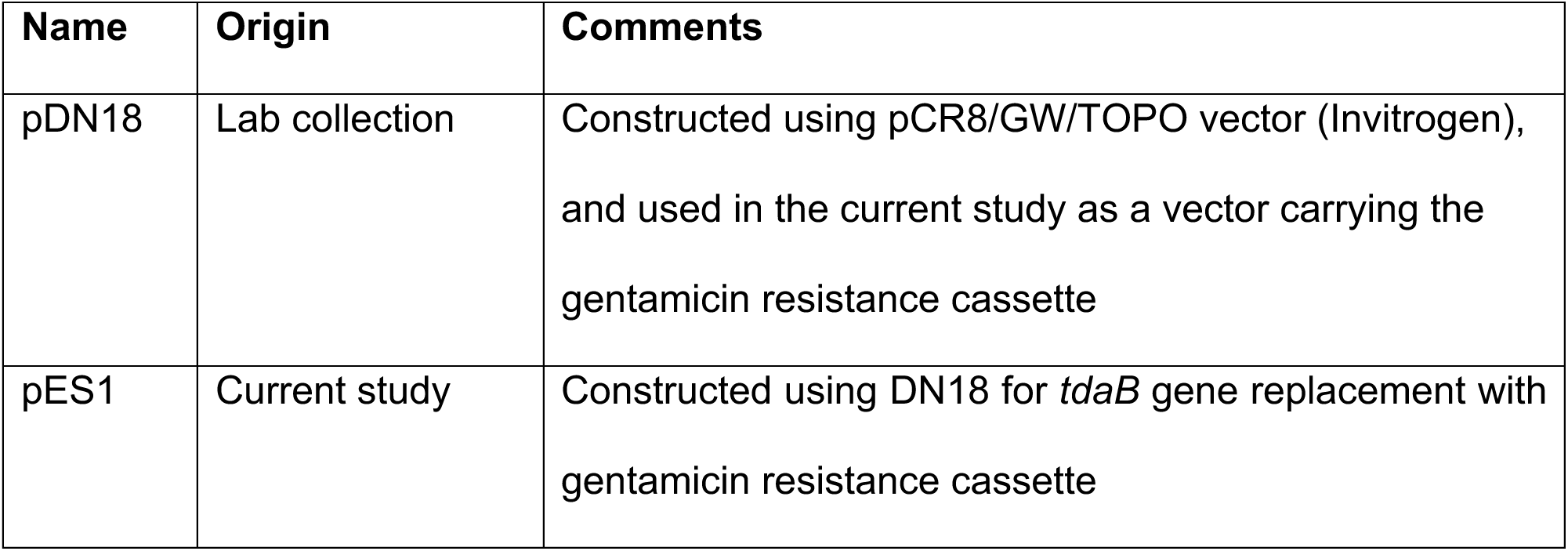
Plasmids used in this study.

Preparations of competent cells was performed as previously described (48), with slight modifications. Briefly, cells were grown to an OD_600_ of 0.7 in ½ YTSS with 40 g/L of sea salts (also termed full salts medium). Bacteria were washed 3 times using 10% (v/v) ice-cold glycerol, centrifuged each time for 5 min at 4 °C and 6000 rpm. The competent cells were subsequently adjusted to an OD_600_ of 20 using 10% (v/v) ice-cold glycerol, frozen in liquid nitrogen and stored at -80°C. Electroporation was conducted with 300 μl aliquots of electrocompetent cells, using 10 µg of DNA in a 2 mm cuvette at 2.5 V, followed by 4 h recovery in ½ YTSS with full sea-salts concentration. The transformed cells were then plated on ½ YTSS plates containing 30 μg/ml gentamicin, and resistant colonies were validated by PCR (Table 2, primers 1025 and 1026) and DNA sequencing.

### Bacterial growth in mono-cultures

Bacteria were grown in a 96-well sterile plate in CNS medium with 5.5 mM succinate as the carbon source. An initial inoculum of OD_600_ = 0.01 was introduced into 150 µl medium in each well, overlaid with 50 µl of hexadecane (Thermo Fisher Scientific, UK) to prevent evaporation. Growth was monitored in 30°C using the Infinite 200 Pro M Plex plate reader (Tecan Group), and measurements were performed each hour after shaking.

To transform the obtained absorbance values into cells/ml, the absorption values were converted to OD_600_ values by multiplying by a conversion factor of 3.8603 as reported previously (1). Then, values were multiplied by a factor of 325,679,613 that was calculated to correspond to cells concentrations (cells/ml) for OD_600_ = 1 for *P. inhibens* WT strain. This factor was obtained as an average of corresponding OD_600_ and CFU/ml values for 10 different time points (using biological duplicates).

### TDA susceptibility plate assay

Liquid pre-cultures of *D. shibae*, *P. inhibens* WT and *P. inhibens* Δ*tdaB* were prepared as described earlier. *D. shibae* was plated using sterile cotton swabs on MB agar plates to create a lawn. The lawn was dried, and on top of it, 5 μl of *P. inhibens* WT or *P. inhibens* Δ*tdaB* cultures (in duplicates) were placed and were allowed to dry. Plates were incubated 5 days at 30°C allowing visible colonies of *P. inhibens* WT or Δ*tdaB* to develop. The pigmentation and the presence of an inhibition zone around the *P. inhibens* WT or Δ*tdaB* colonies were evaluated.

### Co- and tri-culture preparation

Bacteria were grown in pre-cultures as described earlier. For co-cultures, bacteria were inoculated in 10 ml CNS medium with 5.5 mM succinate as the carbon source in final concentration of OD_600_ = 0.005 calculated for each bacterial strain (OD_600_ of pre-cultures were measured with spectrophotometer Ultrospec 2100 Pro, Biochron). Co-cultures were cultivated in 30°C in the dark with shaking (130 rpm).

For tri-cultures, algal cells were inoculated in ASW with L1-Si supplements (250 ml Erlenmeyer flasks) in a final concentration of 10,000 cells per 30 ml of medium. Four days later, bacteria from liquid pre-cultures were diluted in sterile ASW and added to algae to a final concentration of no more than 50 cells/ml for each strain (bacterial concentrations in pre-cultures were estimated by OD_600_ measurements). The final concentration of bacterial cells was confirmed by colony counts on agar plates (as described below). Cultures were kept in 18°C under a light/dark cycle of 16/8 hours (130 μmol photons m^−2^ s^−1^) without shaking. Each time the flask was mixed for the sampling purposes, it was eliminated from the experiment.

### Monitoring bacterial growth in co- and tri-cultures

#### Bacterial enumeration via CFU

To estimate bacterial cell concentrations in the cultures, colonies were counted on selective agar plates. For this purpose, 10 μl of a diluted culture were plated on ½ YTSS (*P. inhibens*), ½ YTSS with 30 μg/ml gentamicin (*P. inhibens* Δ*tdaB*) or ½ MB with 10 μg/ml kanamycin (*D. shibae*) agar plates. After two days for *P. inhibens* and five days for *D. shibae,* colonies were counted and a colony forming units per ml (CFU/ml) value was calculated based on the dilution factor. Two independent technical repeats on separate agar plates per each biological replicate were performed.

#### Bacterial enumeration via qPCR analysis

Bacteria are known to form aggregates when cultivated with algae (2), and this might affect the results of CFU counts on agar plates. Therefore, to validate the bacterial counts in tri-cultures, a quantitative PCR (qPCR) method based on DNA copy number was used as an independent enumeration method.

For this, 20 ml of tri-cultures after 8 days of cultivation (in triplicates) were pelleted by centrifugation at 13,000 g for 2 min at 4°C. Samples were kept on ice at all times to prevent DNA degradation, and the pellets were kept at -20°C until DNA extraction. Genomic DNA was extracted using Wizard Genomic DNA Purification Kit (Promega, USA) following the manufacturer protocol for bacterial DNA extraction. The final concentrations of extracted DNA were measured with Quibit HS dsDNA assay (Thermo Fisher Scientific, USA).

A species-specific set of primers was designed for each bacterial species using the Mauve software (49), BioCyc (50) and NCBI (51) databases (Table 4). The product sizes were 130 bp for all primer pairs. A calibration qPCR confirmed no off-target amplification with both bacterial and algal genomes, for each pair of primers.

**Table 4:**
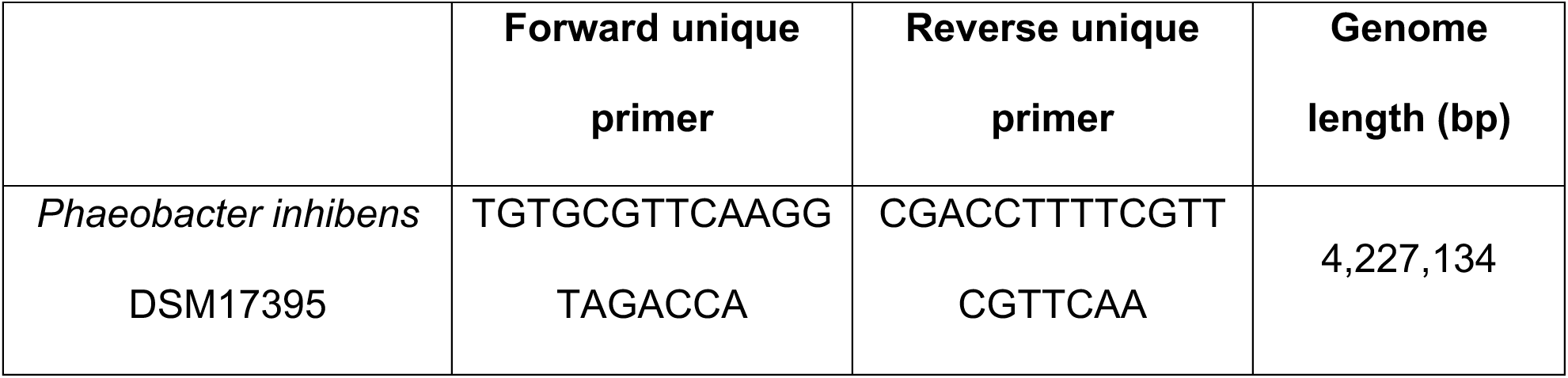
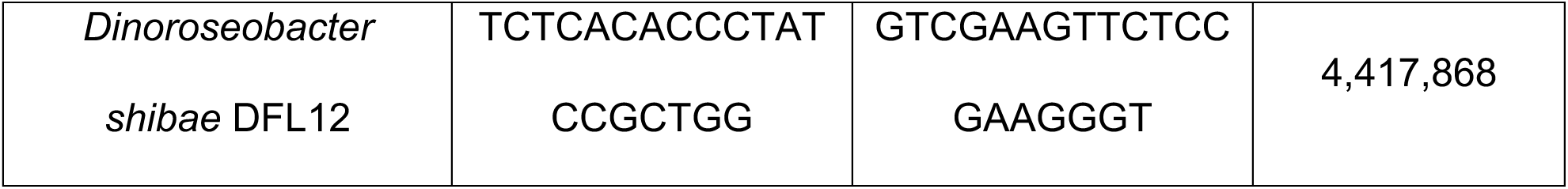
Primers used in this study for qPCR, and bacterial genome length.

qPCR was conducted in 384 well plates, using SYBR Green Master mix (Thermo Fisher Scientific, USA) in a QuantStudio 5 (384-well plate) qPCR cycler (Applied Biosystems, Foster City, USA). The qPCR program ran according to the enzyme requirements for 30 cycles (annealing temperature 60°C). Primer efficiencies and standard curves were obtained by qPCR amplification of known DNA concentrations and analyzed using QuantumStudioTM Design & Analysis Software (Thermo Fisher Scientific, USA). The results of the experiments were analyzed using the obtained standard curves.

To convert ng of DNA into copy numbers of bacterial genomes, the following formula was used, as previously reported (11):

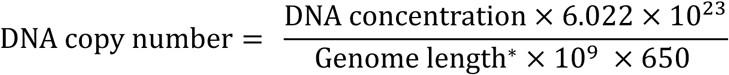

6.022×10^23^ – Avogadro’s number

10^9^ – Conversion from g to ng

650 – Average weight of a bp (g/mole)

* Genome lengths of bacterial strains are listed in Table 4.

### Monitoring algal growth

Algal concentrations in cultures were measured by a CellStream CS-100496 flow cytometer (Merck, Darmstadt, Germany), using 561 nm excitation and 702 nm emission wavelength. All algal samples included three biological replicates. The only exception was the data point for algal growth on day 9 (Figure 4C), which included two biological replicates due to a technical sample loss. For each sample 50,000 events were recorded (with the exception of day 0), and all measurements were performed using technical duplicates. For day 0, due to the dilute nature of the culture, 1,000 event were recorded and the average value of three biological replicates, before bacterial addition, was calculated. Algal cells were gated according to event size and fluorescence intensity.

### Statistics

At least 3 biological replicates were used for each treatment. For each biological replicate, the mean of at least 2 technical repeats was calculated. Statistical analysis included two-tailed t-test and was performed using Spyder 6.0.1 (Python Software Foundation).

## Acknowledgments

We are grateful to Dr. Jorge Rocha from the Centro de Investigaciones Biológicas del Noroeste, Mexico, for his invaluable assistance in the initial stages of genetic engineering in marine bacteria. We thank all members of the Segev lab for their insights and contributions during our discussions.

This study was supported by funds received from the European Research Council (ERC StG 101075514), the Israeli Science Foundation (ISF 692/24), and the de Botton Center for Marine Science, granted to E.S.

## Data availability

All data are available in the article or Supplementary Information. Plasmids and bacterial strains generated in this study will be made available on reasonable request.

